# Eyes robustly blink to musical beats like tapping

**DOI:** 10.1101/2024.07.04.602077

**Authors:** Yiyang Wu, Xiangbin Teng, Yi Du

## Abstract

Auditory-motor synchronization with musical rhythm may extend beyond overt body movements like fingers or feet. Through four experiments combining eye-tracking, neurophysiological and structural imaging approaches with 123 young non-musicians, we demonstrated robust synchronization of spontaneous eye blinks with musical beats. The blinking rate consistently aligned with the beat rate across various tempi and independent of melodic cues. Blinking exhibited beat phase-specific tuning, with inhibition immediately before beats and increased frequency afterward. Variations in blink-beat synchronization corresponded to the difference in microstructural lateralization of auditory-parietal connectivity. EEG recordings revealed a dynamic correspondence between blink and neural beat tracking. Mechanistically, this synchronization reflects dynamic auditory attention and temporal modulation of visual sampling. Our findings establish ‘eye tapping’ as a novel behavioral paradigm, expanding the behavioral repertoire of auditory-motor synchronization. It underscores the intricate relationship between music rhythms and peripheral oculomotor system, proposing a cross-modal active sensing and embodied experience in music perception.

## Introduction

Auditory-motor synchronization—voluntarily coordinating our body movements with the rhythmic patterns of music, such as tapping our feet or fingers to the beat—is a universally observed human behavior(*1*). This phenomenon, which is not as prevalent among species closely related to humans, points to the evolution of specialized auditory-motor neural circuitry within the human brain(*2–5*). Delving into this synchronizing behavior not only advances our understanding of its distinct neural circuitry but also provides insights into brain diseases, as rhythm and timing have been identified as vulnerabilities in neurodevelopmental disorders like autism spectrum disorder and developmental language disorder(*6*). Despite extensive studies, however, the full diversity of such synchronizing behaviors remains incompletely characterized. In this work, we unveil a previously unrecognized form of synchronization: eye blinks consistently aligning with musical beats. This discovery suggests a novel, functional, and neural linkage between cortical auditory processing and peripheral oculomotor mechanisms.

Enhanced comprehension of behaviors often heralds deeper insights into neural systems(*7–9*). Observations of synchronized behaviors, such as voluntary tapping, dancing, and nodding, have spurred theories that auditory-motor interactions predominantly engage the central nervous system(*10–13*). A notable recent model suggests a pathway from auditory regions to subcortical areas, wherein rhythmic acoustic patterns entrain the basal ganglia and cerebellum, resulting in motor activities that synchronize with sound patterns(*2, 14–17*). Alternatively, some research points to a cortical pathway for beat tracking, involving connections between the premotor and auditory cortical areas(*18–22*). Although these discoveries have greatly advanced our understanding of auditory-motor synchronization, they also raise the question: are the behaviors these models are predicated upon fully representative?

The influence of music on the brain is profound and complex(*23*). It is plausible that music-induced synchronized behaviors extend beyond overt physical movements, such as finger tapping and dancing, to more subtle, involuntary actions. Music, with its rhythms and pitches, can evoke intense emotional responses, often described as ‘chills’, and activate the dopamine system, thereby influencing overall brain states(*24, 25*). Given the intricate connections between peripheral or autonomic systems and both emotional states and dopamine pathways, music could modulate and even entrain automatic physiological responses. This prompts an intriguing line of inquiry: how extensively can music synchronize with not only overt physical movements but also with spontaneous autonomic system-governed activities? This study investigates the synchronization of musical rhythms with eye movements, specifically the spontaneous oculomotor activities that are part of the peripheral or automatic systems(*26, 27*).

Previous studies have explored the intricate relationship between oculomotor activity and music listening experiences(*28, 29*), placing a significant focus on pupillary responses. Variations in pupil size have been linked to cognitive and emotional states(*30–34*). While attempts have been made to associate pupillary activity with musical rhythms, they primarily concentrated on detecting unexpected patterns(*35–37*), linking pupil dynamics to temporal attention and specific note configurations(*38*), and more recently, directly to musical beat structure(*39*). Apart from pupil size, research has demonstrated that eye blinks can track artificial sentence structures, somewhat resembling high-level musical structures, though not musical beats(*40*). However, the connection between eye movements and music’s rhythmic beats remains unestablished.

Our objective is to delve into the underexplored connection between peripheral oculomotor activity, specifically eye blinks, and fundamental musical rhythms such as beats. We aim to provide behavioral evidence, unveil the neural and structural substrates involved, and uncover the underlying mechanism. To achieve this, we first exposed participants to Western classical music characterized by highly regular beat patterns, while monitoring their ocular activities. Additionally, we recorded neural responses to examine how musical structures are encoded in the brain and their correlation with ocular activities. Furthermore, we assessed the microstructural properties of white matter tracts connecting frontal, parietal and auditory regions to investigate the structural foundation of individual variations in auditory-oculomotor synchronization. Lastly, we explored the functional significance of this synchronization between eye movements and music. Overall, our research reveals a robust new phenomenon of auditory-motor synchronization in music listening: ‘eye tapping,’ and offers a mechanistic explanation by integrating behavioral, neural, and structural evidence.

## Results

### ‘Eye tapping’: Eye blinks reliably track musical beats

In Experiment 1, 30 non-musicians listened to 10 Bach chorales at 85 beats per minute in both the original and reverse versions, while undergoing electroencephalogram (EEG) and eye-tracking recordings simultaneously. In the reverse version, the sequence of beats was inverted from end to beginning while maintaining temporal regularity at the beat level and the original’s acoustic properties (**Fig. 1B and C**). This design ensured the reverse version remained an unfamiliar control. Each musical piece was repeated three times consecutively to collect sufficient data on a single piece and measure the effect of repetition. The experimental procedure is summarized in **Fig. 1A and B**. The acoustic modulation spectra of all musical pieces are shown in **Fig. 1C**, demonstrating prominent frequency components of musical beat structures (beat rate: 1.416 Hz). The liking rating showed no difference between the original and reverse versions (**Fig. 1D**).

**Fig. 1.**
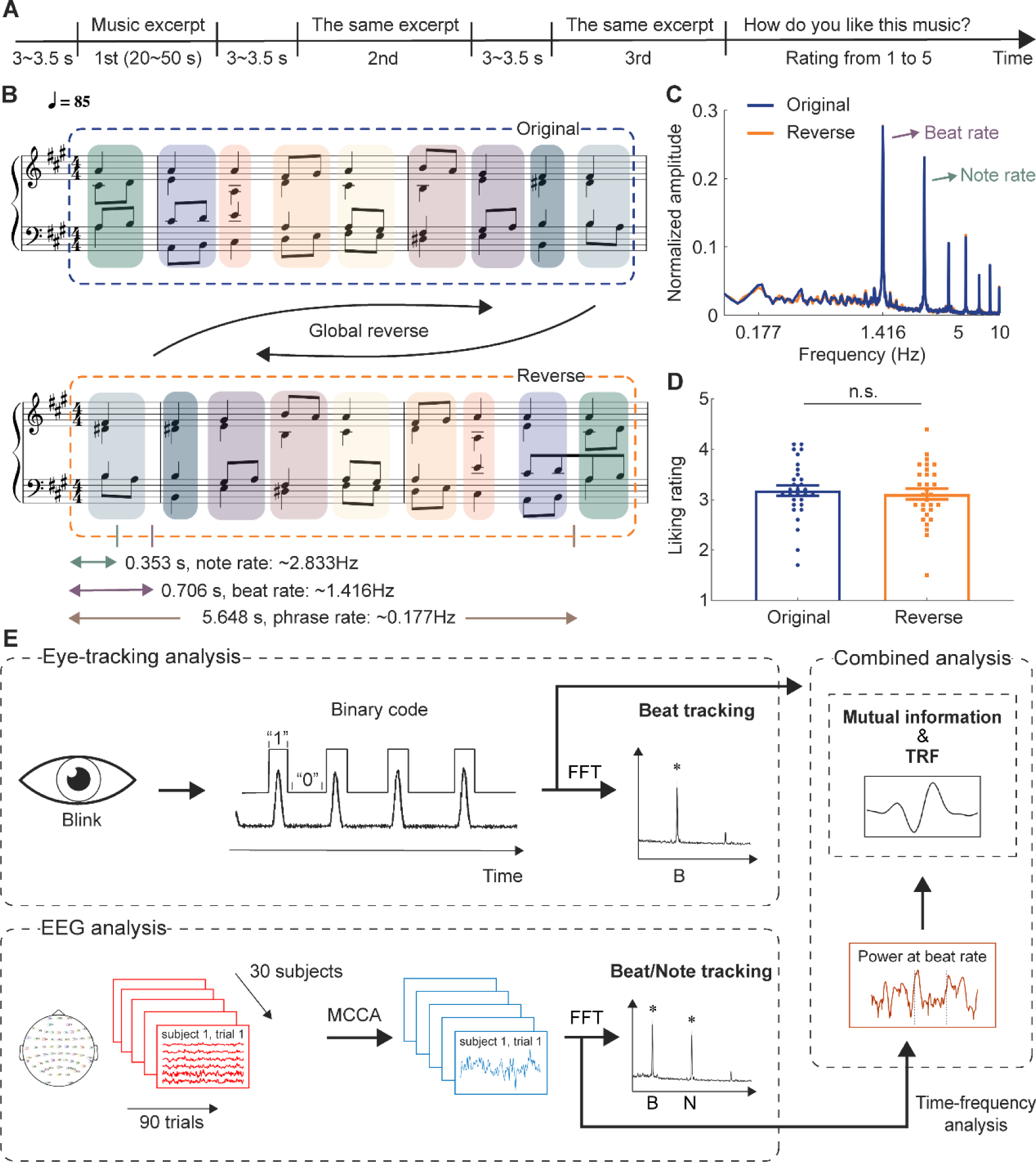
Design and data analysis pipelines of Experiment 1. (**A**) Each music excerpt was presented three times consecutively undergoing EEG and eye-tracking recordings. After each trial, participants rated how much they liked the excerpt. (**B**) Example of music excerpts. The tempo is 85 beats per minute: each note is ∼353 ms (2.833 Hz), each beat is ∼706 ms (1.416 Hz), and each phrase is ∼5,648 ms (0.177 Hz). The reverse version (bottom) was created by reversing the original excerpts beat order (top). (**C**) Modulation spectra of the stimuli showed peaks at the note, beat rates and harmonics for both versions. (**D**) Liking rate showed no significant difference between the versions. (**E**) Overview of the analysis. Blinks were recorded via an eye tracker and converted into binary strings. Beat tracking was measured by the fast Fourier transform (FFT) of the blink signals. EEG was recorded with 64 electrodes, and multiway canonical correlation analysis (MCCA) extracted the most common component across subjects. Denoised signals measured neural tracking of beats and notes. Neural signal power at the beat rate was obtained using a wavelet transform. Mutual information and temporal response function (TRF) measured the relationship between blink and neural signals.

Next, we measured the neural tracking of musical beats, as well as eye-blink tracking of beats. We extracted music-related EEG components (see Methods) and applied the fast Fourier transform to neural signals, and derived amplitude spectra to examine whether there were neural components that corresponded to musical structures (**Fig. 1E**), a routine analysis of neural tracking of musical structures. In a similar vein, we quantified the continuous eye-tracking data to extract the timing of eye blinks (by converting the data to binary time series) and then derived the amplitude spectra of eye blinking (**Fig. 1E**).

To our surprise, while the neural tracking of musical beats has been well documented(*41–45*), a similar finding was shown in the spectra of eye blinking; that is, eye blinks track musical beat structure—eye tapping (**Fig. 2**). The eye tracking of musical beats was significant, as shown in **Fig. 2A**, demonstrated by the fact that the spectral peaks of eye blinking dynamics were above the thresholds (*p* < 0.01) created through a surrogate test on the group-averaged data (see Methods section for details). We further derived a significant frequency range for further analyses: the frequency range was from 1.4 Hz to 1.4333 Hz (see Methods for details).

**Fig. 2.**
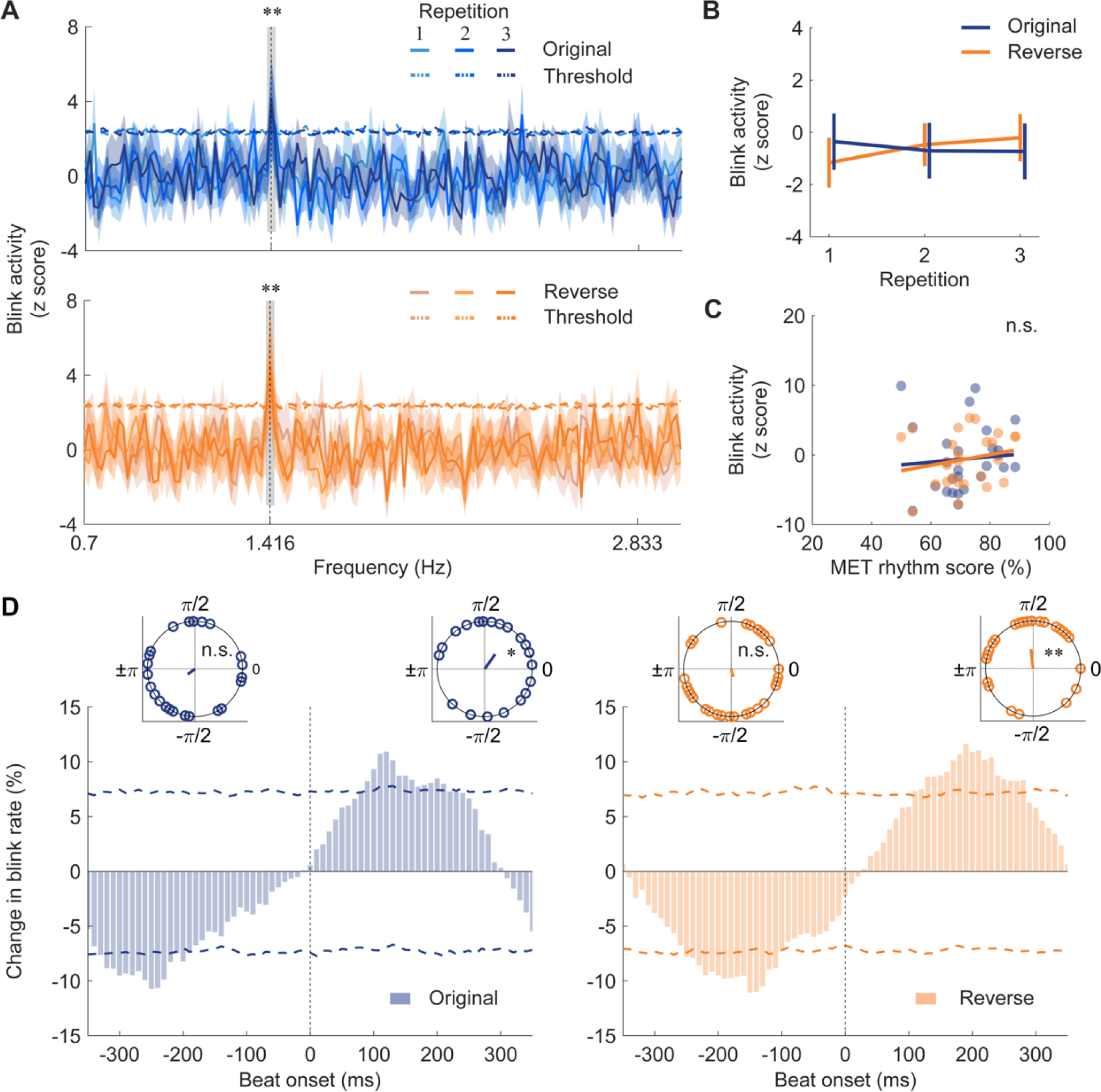
Blink tracking of musical beats. (**A**) Blink amplitude spectra for two versions and repetitions, with horizontal dashed lines showing the threshold from the surrogate test (*p* < 0.01). Shaded areas represent 1 SEM across participants (n = 30). Gray boxes indicate frequency ranges where amplitude was above the threshold. The blink activity showed a salient peak around the beat rate. (**B**) Blink response amplitude within significant frequency ranges around the beat rate, with error bars denoting 1 SEM. Version and repetition had no significant effect on blink response. (**C**) Correlation between the MET rhythm score and blink response amplitude, showing no significant relation. Colored dots represent individuals (n.s., not significant). (**D**) Blink rate during listening, with histograms showing percent change relative to the mean permuted data. Vertical dashed lines mark beat onset. Horizontal dashed lines represent 0.01 and 0.99 levels for blink rate change. Inset phase plots show individual phase angles of the highest (right) and lowest (left) values relative to beat onset

The finding, though surprising and perhaps entirely new, is robustly observed across the two versions of musical pieces and the three repetitions. This is further echoed by a histogram of raw intervals between two consecutive blinks, as shown in **Supplementary Fig. 1**, with a clean peak of intervals between one and two beats. In the following analyses, as neural tracking of beats has been demonstrated and not our current focus, we focused on the eye blink tracking of beats and its relationship with neural signals. Given the normal spontaneous blink rate is between 12 and 15 per minute(*46*), the eye tapping rate we showed here is highly specific to the musical tempo used.

To assess the effects of different versions of musical pieces and repetition times on eye tracking, we conducted a two-way repeated-measures ANOVA with Version (Original or Reverse) and Repetition (1, 2 or 3) as main factors on corrected amplitude within significant frequency range (**Fig. 2B**). We did not find any significant main effect nor interaction effect (version: *F*_(1,25)_ = 0.001, *p* = 0.973, *η_p_^2^* = 0.000; repetition: *F* _(2,50)_ = 0.117, *p* = 0.890, *η_p_^2^* = 0.005; interaction: *F*_(2,50)_ = 0.642, *p* = 0.531, *η_p_^2^* = 0.025). The reason for this result could be that as beat structures are well preserved in different versions of musical pieces and are regular to follow, participants can track the beat structures easily in each condition. As a result, even though the eye tracking of beats is

We then tested whether the eye tracking correlates with musical ability measured by the rhythm subtest of the Musical Ear Test (MET)(*47*). The corrected amplitude within significant rhythm score to examine the effect of musical rhythmic ability. We didn’t find any statistical significance for each version (original: *r*_(26)_ = 0.241, *p* = 0.236; reverse: *r*_(26)_ = 0.240, *p* = 0.238; **Fig. 2C**). No significant correlation was found probably because, as stated above, the beat structure is easy to track in our stimuli.

This phenomenon, the eye tracking of beats, is akin to finger tapping to musical beats, so we termed this music-oculomotor behavior as ‘eye tapping’.

### Temporal signature of eye tapping shows phase selectivity

We next investigated how eye blinks ‘tap to’ musical beats from a temporal perspective. The hypothesis here is that, like finger tapping(*12, 13*), eye blinks should center around certain phases of a beat during eye tapping. If the eye blinks tracking musical beats are indeed responses to auditory stimulation (i.e., musical notes), it would be expected that in line with recent evidence of eye blinks induced by auditory and visual stimulation, the latency of eye blinks to the beat onset would predominantly center around 500 milliseconds (ms) or after(*40, 48*). Conversely, if the eye blinks are simply the corneal reflex or blinking reflex caused by the sudden onset of note sounds, the latency would average around 60 ms(*49*).

**Figure 2D** shows the normalized timing distribution of eye blinks, or the eye blinking rate, referenced to each musical beat onset (see Methods). Following procedures in previous studies(*50*), a permutation test was conducted and revealed that eye blinks significantly decreased before the beat onset, referred to as ‘blink inhibition’ in the present study, and increased starting around 100 ms after the beat onset. Circular statistic conducted with V-test showed that eye blinks were time-aligned to the post-beat period (original: *v* = 9.418, *p* = 0.008; reverse: *v* = 12.37, *p* = 0.001; V test for non-uniformity with π/2), while blink inhibition occurred randomly during pre-beat period (original: *v* = 5.566, *p* = 0.075, V test with −3π/4; reverse: *v* = 5.032, *p* = 0.097, V test with −π/2). This finding suggests that the temporal range of eye blink latencies unveiled here extends beyond the corneal reflex but precedes the typical 500 ms latency shown in sensory stimulation-induced eye blinks.

### Eye tapping corresponds to neural tracking of musical beats

As neurophysiological signals were also recorded in Experiment 1, we first replicated the established phenomenon of neural beat tracking. Subsequently, we examined the correspondence between eye tracking and neural tracking of beats. The eye-neural correspondence could potentially elucidate what led eye blinks to track musical beats.

In line with previous studies(*41–45*), robust neural tracking of beats was evident at the beat rate (**Fig. 3A**). After preprocessing raw EEG data and removing independent components associated with eye blinks, eye movements, and heartbeat using an independent component analysis (ICA) algorithm, we applied multiway canonical correlation analysis (MCCA) to extract components linked to music listening (see Methods). This approach enabled the extraction of auditory neural signals specific to music listening and the analysis of single-trial data for each musical piece and repetition. We then derived power spectra of neural signals for music listening, normalized the magnitude, extracted the corrected amplitude within the significant frequency ranges (at beat rate, 1.3833 – 1.45 Hz; at note rate, 2.8 – 2.9 Hz) for both beat and note rates, and conducted the permutation test.

**Fig. 3.**
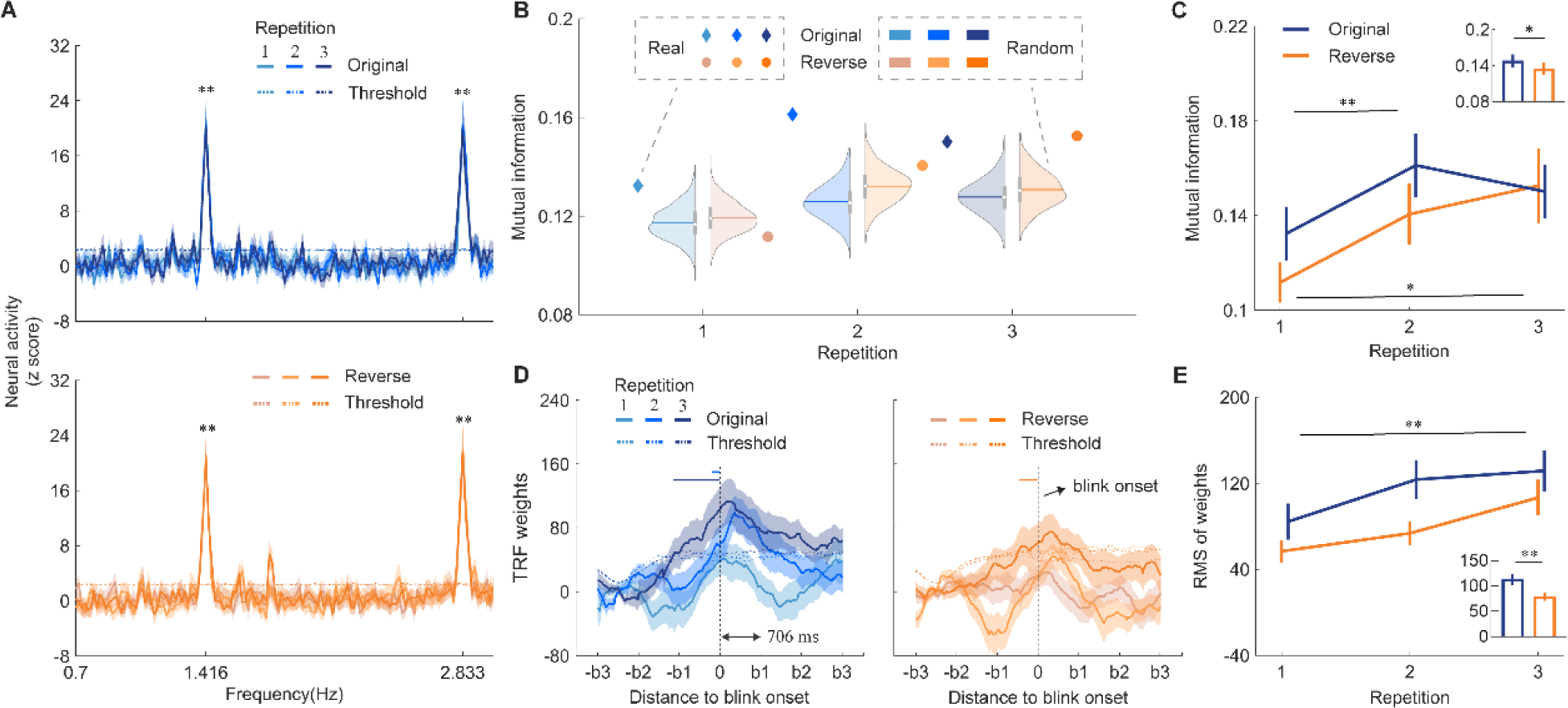
Combined analysis on blink and neural signals. (**A**) Neural response spectra for two versions, with shaded areas representing 1 SEM (n = 30). Horizontal dashed lines represent surrogate test thresholds (*p* < 0.01). Salient peaks appeared at the note and beat rates for both versions. (**B**) Mutual information (MI) between blink signals and beat-rate EEG power. Violin plots display MI distribution between blink signals and circular shifted EEG data. (**C**) Averaged MI for each version and repetition. Bar plot shows mean MI values for both versions. Error bars denote 1 SEM. MI increased with repetition and was higher in the original version. (**D**) Temporal response function (TRF) of beat-rate EEG power using blink onset as the regressor. Vertical dashed lines mark blink onset. Horizontal axis covers three beats before and after blink onset. Horizontal dashed lines show permutation test thresholds (*p* < 0.01). Notably, TRF weights rose robustly before blink onset, as shown by horizontal solid lines. (**E**) Root mean square (RMS) of TRF weights at peak point. Bar plot shows mean TRF weights for both versions. Error bars denote 1 SEM. RMS of weights was larger in the original version. * *p* < 0.05, ** *p* < 0.01.

To examine the effects of reversal manipulation and repetitions, we performed a two-way repeated-measures ANOVA of Version and Repetition for beat and note rates separately (**Supplementary Fig. 2**). At the note rate, the analysis showed no main effect of Version (*F*_(1,26)_ = 0.948, *p* = 0.339, *η_p_^2^* = 0.035) nor interaction (*F*_(2,52)_ = 0.718, *p* = 0.493, *η_p_^2^* = 0.027), but a main effect of Repetition (*F*_(2,52)_ = 4.078, *p* = 0.023, *η_p_^2^* = 0.136). At the beat rate, neural tracking of beats was not significantly modulated by either factor or their interaction (version: *F*_(1,27)_ = 0.196, *p* = 0.661, *η_p_^2^* = 0.007; repetition: *F*_(2,54)_ = 1.566, *p* = 0.218, *η_p_^2^* = 0.055; interaction: *F*_(2,54)_ = 1.334, *p* = 0.272, *η_p_^2^* = 0.047). Additionally, neural tracking of notes and beats correlated with the MET rhythm score solely for the reverse version (note rate: *r*_(27)_ = 0.445, *p* = 0.020; beat rate: *r*_(28)_ = 0.454, *p* = 0.015), suggesting that a higher level of musical ability is necessary for effectively tracking beat structures in the reverse version. The potential explanation could be that the harmonic progressions in the reverse versions are unusual to listeners. Consequently, this demands enhanced rhythmic skills to accurately follow the music’s beat patterns upon first exposure to these reverse versions of musical pieces.

Having observed both eye tracking and neural tracking of musical beats, we now examined their relationship through mutual information (MI) and eye-blink triggered temporal response function (TRF). MI between eye blink timing series and neural signals of beat tracking across entire musical pieces can indicate whether the correspondence exists. Indeed, compared to the surrogate data, empirical MI values were statistically significant for all repetitions in the original version (*ps* < 0.05; **Fig. 3B**). However, for the reverse version, a significant result was observed only on the third presentation (*p* = 0.905, *p* = 0.12 and *p* = 0.002; first, second and third presentations respectively). A two-way repeated-measures ANOVA revealed a significant main effect of Version (*F*_(1,27)_ = 4.396, *p* = 0.046, *η_p_^2^*= 0.140; **Fig. 3C**), with higher MI in the original version than the reverse. It also revealed a significant main effect of Repetition (*F*_(1.629,43.979)_ = 9.767, *p* = 0.001, *η_p_^2^* = 0.266), with higher MI on the second and third presentations than the first (*p* = 0.002 and *p* = 0.010, second and third presentations respectively). The interaction was not significant (*F*_(2,54)_ = 1.479, *p* = 0.237, *η_p_^2^* = 0.052).

Interestingly, MI results demonstrated that repetitions or listeners’ familiarity with musical pieces modulated the correspondence between eye tracking and neural tracking of beats, despite no such effect being observed in either neural tracking (**Supplementary Fig. 2**) or eye tracking (**Fig. 2A and B**), individually. The significant correspondence was evident across all repetitions of the original version but only emerged during the third repetition of the reverse version. This discrepancy between the original and reverse versions, despite featuring identical beat structures, implies a more effective anticipation facilitated by normal harmonic progressions in the original version in binding eye tracking and neural tracking. Furthermore, the repetition effect again underscores that eye tapping transcends mere reflexive responses to sensory stimulation.

### Neural dynamics of beat tracking precede eye tapping

Next, we investigated the temporal relationship between eye tapping and neural tracking. We aimed to assess whether the dynamics of one index precede, if not predict, the other. Employing TRF methodology, we treated eye blinks as events or features and examined the fluctuations in neural power of beat tracking before and after eye blinks.

Remarkably, we observed an increase in neural power of beat tracking, as reflected in TRF weights, before the onset of blink (**Fig. 3D**). Specifically, for the original version, the TRF weights were significantly higher than threshold derived from surrogate data before blink onset during the second and third presentations. Similarly, for the reverse version, TRF weights exceeded threshold before blink onset during the third presentation. The repetition effect observed here echoes the MI results. A two-way repeated-measures ANOVA on the root mean square (RMS) of TRF weights at the response peak (see Methods) found a significant main effect of Version (*F*_(1,27)_ = 10.808, *p* = 0.003, *η_p_^2^* = 0.286; **Fig. 3E**), suggesting higher RMS of weights for the original version than the reverse version. It also showed a significant main effect of Repetition (*F*_(2,54)_ = 5.150, *p* = 0.009, *η_p_^2^* = 0.160), with higher weights during the third presentation than the first presentation (*p* = 0.002, Bonferroni corrected), indicating a gradual formation of prediction for blink onset in the brain.

Thus far, we have established that eye tapping—wherein eye blinks track musical beats— corresponds robustly with neural tracking of beats. Moreover, eye tapping transcends mere passive reflex to sensory stimulation, and the eye-neural correspondence is modulated by cognitive factors such as repetition and reversal manipulations.

### Structural basis of individual differences in eye tapping

Previous studies have revealed notable individual differences in people’s abilities to synchronize with auditory-motor activities, such as whispering or clapping in synchrony with auditory stimuli(*51, 52*). This individual variability was also evident in our study: after excluding 4 outliers, a k-means clustering algorithm utilizing blink response amplitude partitioned the data into two distinct clusters (**Fig. 4A**). The corrected amplitude of blink tracking was higher for the second cluster (high synchronizers, n = 12) compared to the first cluster (low synchronizers, n = 14). However, no significant difference was observed between the two groups concerning their MET rhythm score (*t*_(24)_ = −1.211, *p* = 0.238, Cohen’s *d* = −0.481; **Fig. 4B**).

**Fig. 4.**
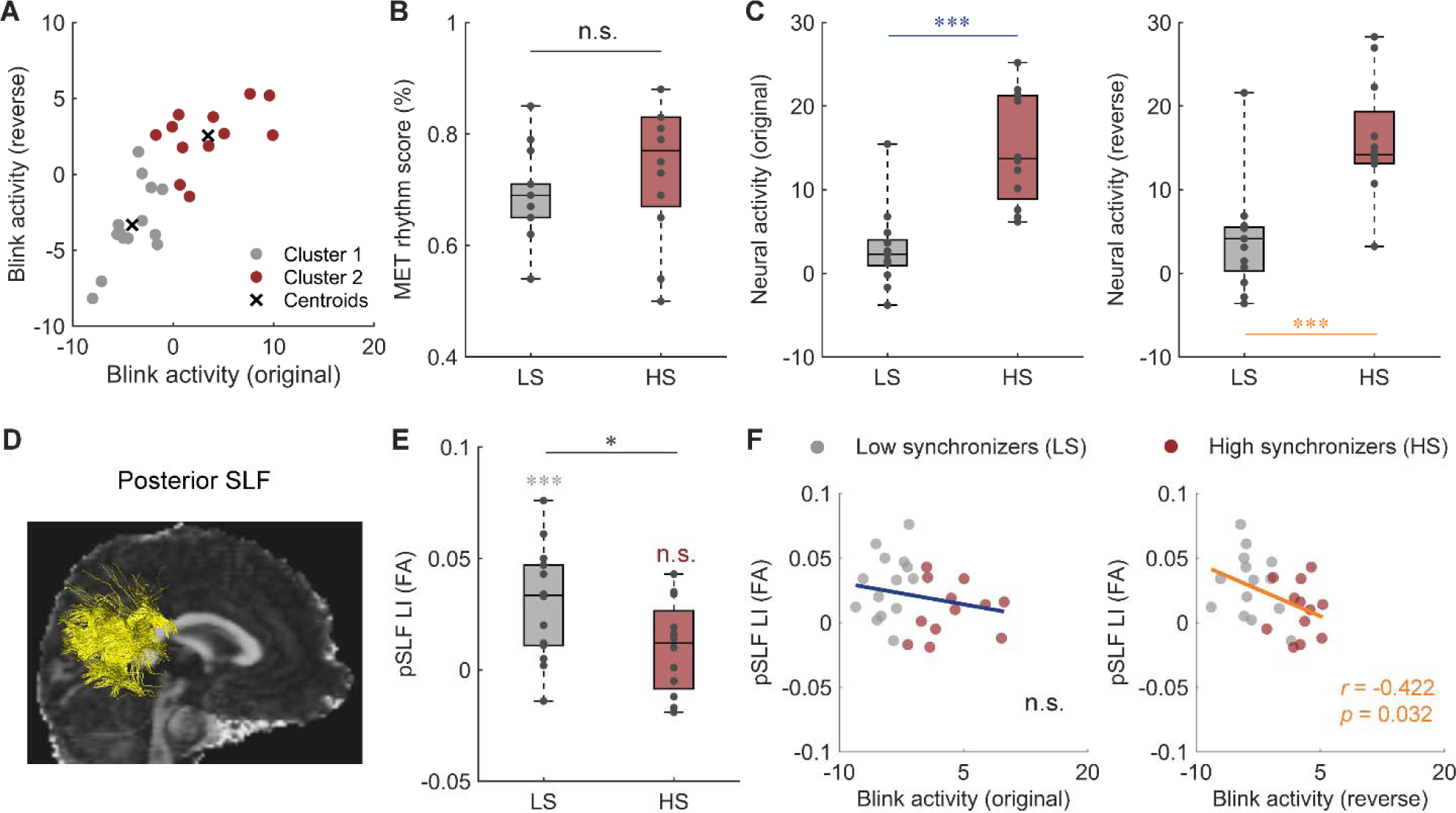
Individual differences in eye tapping and microstructural properties. (**A**) Cluster assignments and centroids (black crosses) of the blink response amplitude data. Gray dots correspond to low synchronizers (LS, n = 14) and red dots correspond to high synchronizers (HS, n = 12). The results showed a group segregation of eye tapping ability. (**B**) No significant difference in musical ability (MET rhythm score) between the two groups. (**C**) EEG response amplitudes at beat rate (neural tracking of beat) was stronger in high synchronizers than in low synchronizers for both the original (left) and reverse (right) versions. (**D**) The tractography of right posterior superior longitudinal fasciculus (pSLF) in one typical participant. (**E**) The fractional anisotropy (FA)-based laterality index (LI) of pSLF exhibited greater left lateralization in low synchronizers than in high synchronizers. pSLF was significant left-lateralized in low synchronizers but symmetrical in high synchronizers. (**F**) Correlations between FA-based LI of pSLF and blink activity in the original and reverse versions. Blink activity was negatively correlated with FA-based LI of pSLF for the reverse version. * *p* < 0.05, *** *p* < 0.001, n.s., not significant.

Notably, participants categorized as high synchronizers based on blink response amplitude also showed stronger neural tracking at the beat rate than those categorized as low synchronizers across both versions (original: *u* = 147, *p* = 0.000; reverse: *u* = 141, *p* = 0.000; **Fig. 4C**). Additionally, blink response amplitude correlated with the neural tracking of beats for both versions (original: *r*_(26)_ = 0.643, *p* = 0.000; reverse: *r*_(26)_ = 0.601, *p* = 0.001; Spearman correlation). These findings indicate a robust relationship between neural activities and blink responses, suggesting that the neural response differences may be directly linked to variations in eye tapping. Subsequently, we acquired diffusion-weighted imaging (DWI) data to quantify potential differences in white matter tracts connecting frontal, parietal and auditory regions that might distinguish the two groups in terms of eye tapping ability. Specifically, we focused on two segments of the superior longitudinal fasciculus (SLF), the dorsal SLF (dSLF) and posterior SLF (pSLF), that link auditory and dorsal premotor regions via parietal region (see Fiber tractography in Materials and Methods) and have been proposed to play an important role in auditory-motor synchronization and beat perception(*20, 51, 53*). Microstructural indices (fractional anisotropy (FA), neurite density index (NDI), and orientation dispersion index (ODI)) and their lateralization indices (LI) of bilateral dSLF and pSLF were extracted to explore the relationship between blink data and white matter properties.

Independent t-tests on each index and its LI revealed a significant group difference only in the FA-based LI of pSLF, with low synchronizers showing stronger left lateralization compared to high synchronizers (*t*_(24)_ = 2.156, *p* = 0.041, Cohen’s *d* = 0.848, **Fig. 4E**). Specifically, the pSLF was significantly left lateralized in low synchronizers (*t*_(13)_ = 4.447, *p* = 0.001, Cohen’s *d* = 1.189), whereas no asymmetry was found in high synchronizers (*t*_(11)_ = 1.654, *p* = 0.126, Cohen’s *d* = 0.477). Moreover, the blink response amplitude was negatively correlated with the FA-based LI of pSLF for the reverse version (*r*_(26)_ = −0.422, *p* = 0.032; **Fig. 4F**), implying that a more bilaterally symmetrical or even right-lateralized pSLF may support stronger synchronization of eye blinks to musical beat. No significant group differences were observed on other microstructural indexes (all statistics shown in **Supplementary Table 1**).

### Eye tapping is primarily entrained by the temporal patterns in music

Despite our confirmation of this novel behavior, several questions remain, particularly regarding the elements in musical pieces that induce this behavior. Does the pitch information or melodic patterns crucially drive eye tapping, or can eye tapping be induced solely by temporal patterns in tone sequences deprived of melodic cues? Additionally, given that the eye blink rate during rest falls between 12 – 20 times per minute, considerably slower than the tempi of most musical pieces, does musical tempo modulate eye tapping?

To address these questions, in Experiment 2, we compared eye tapping responses to music and static tone sequences across three tempi (66, 85, and 120 beats per minute) (**Fig. 5A**). Specifically, we generated ten new tone sequences maintaining the temporal structures of the original musical pieces but lacking pitch information or melodic cues. Similar to Experiment 1, 30 non-musicians were instructed to listen attentively to each auditory stimulus while their eye movements were recorded.

**Fig. 5.**
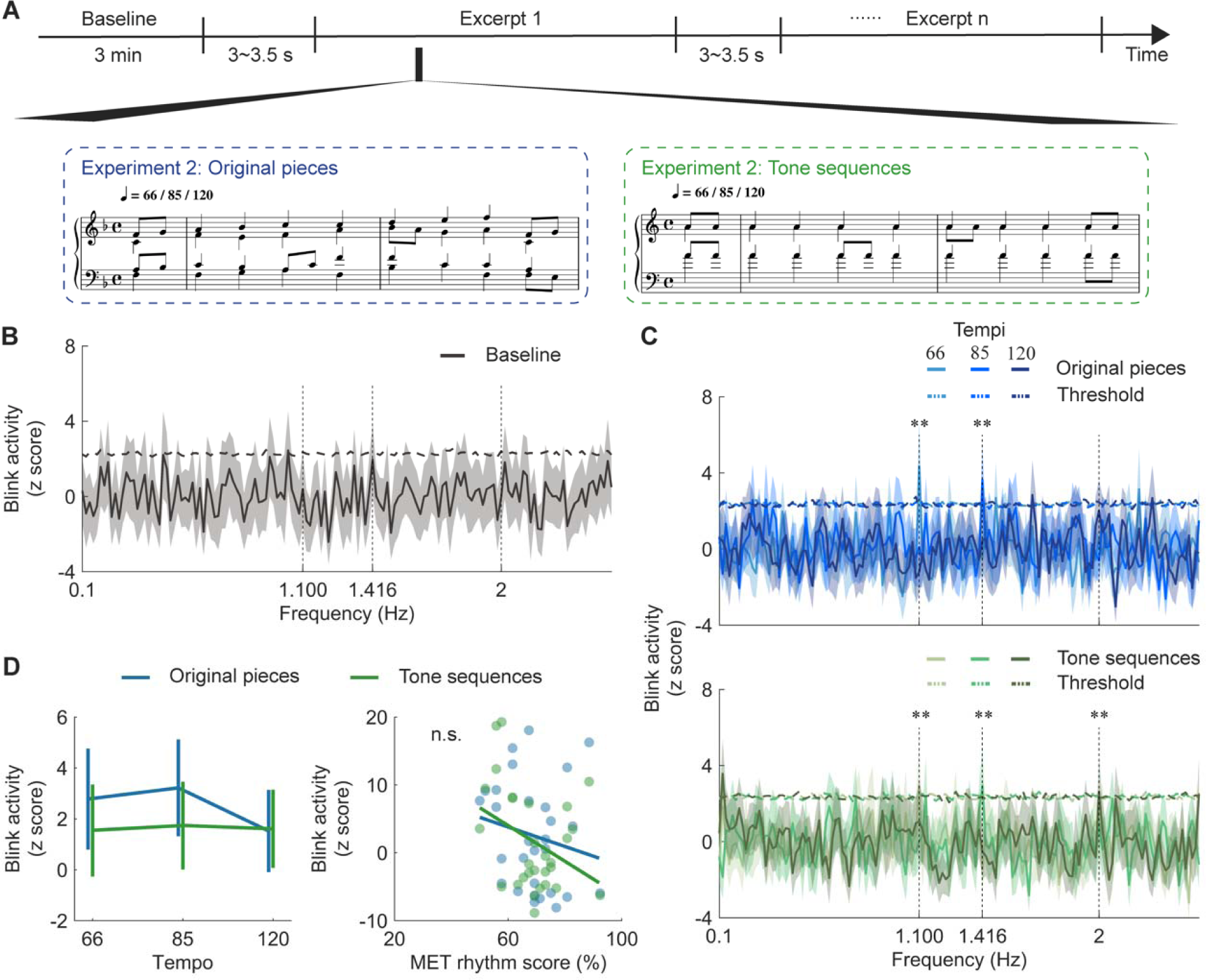
Blink tracking is robust to rhythmic sequences. (**A**) Experimental paradigm for Experiment 2: participants listened to original musical pieces and tone sequences while their eye movements were recorded. 0.01). Shaded areas represent 1 SEM across participants (n = 30). (**C**) Blink amplitude spectra for two stimuli types, with horizontal dashed lines showing surrogate test threshold (*p* < 0.01). Vertical dashed lines indicate frequency points above the threshold. Shaded areas represent 1 SEM. Blink rate showed a salient peak at the beat rate across stimuli types and tempi, except at 120 beats per minute for original pieces. **, *p* < 0.01 by surrogate test. (**D**) Left: Blink response amplitude at the beat rate. Error bars denote 1 SEM. Stimuli type and tempo had no significant effect on blink responses. Right: Correlation between MET rhythm score and blink response amplitude, with no significant relation. Colored dots indicate individuals (n.s., not significant).

One difference from Experiment 1 was the inclusion of a 3-minute rest period before the main experiment to determine whether eye blinks during idle state exhibit any temporal structures. A surrogate test on the baseline data yielded no significant peak (**Fig. 5B**), confirming that the eye tracking peak observed in Experiment 1 was indeed music-related or at least stimulus-related.

In **Fig. 5C**, we replicated our observation in Experiment 1 and noted robust eye tapping in Experiment 2, alongside a modulation effect of musical tempo. We tested the effects of Stimuli type (original pieces or tone sequences) and Tempo (66, 85 or 120 beats per minute) using a two-way repeated-measures ANOVA. Interestingly, though eye tapping appeared to be stronger for original pieces than tone sequences and decreased with increasing tempi, we found no significant main effect (stimuli type: *F*_(1,28)_ = 0.510, *p* = 0.481, *η_p_^2^* = 0.018; tempo: *F*_(2,56)_ = 0.352, *p* = 0.705, *η_p_^2^* = 0.012) or interaction effect (*F*_(2,56)_ = 0.182, *p* = 0.834, *η_p_^2^* = 0.006). Moreover, there was no significant correlation between blink activity and the MET rhythm score (original pieces: *r*_(29)_ = −0.287, *p* = 0.131; tone sequences: *r*_(29)_ = −0.337, *p* = 0.073; **Fig. 5D**).

Taken together, results in Experiments 1 and 2 suggest that eye tapping is primarily entrained by the temporal patterns in music. In other words, pitch changes or melodies do not serve as the main cue for driving eye tapping; instead, eye blinks align with the timing of auditory events.

### Eye tapping facilitates pitch deviant detection

In Experiment 3, we investigated whether eye tapping correlates with cognitive performance. While prior experiments have established a connection between eye tracking and neural tracking of beats, we further sought to determine if better eye tapping corresponds to improved tracking of musical elements? To do so, we conducted a pitch deviant detection task where non-musicians (n = 31) listened to music segments selected from the original musical pieces, each containing two phrases (16 beats in total). A pitch deviant tone was introduced in the second phrase, and participants need to detect deviant tones as soon as possible, while their eye activities were recorded (**Fig. 6A**).

**Fig. 6.**
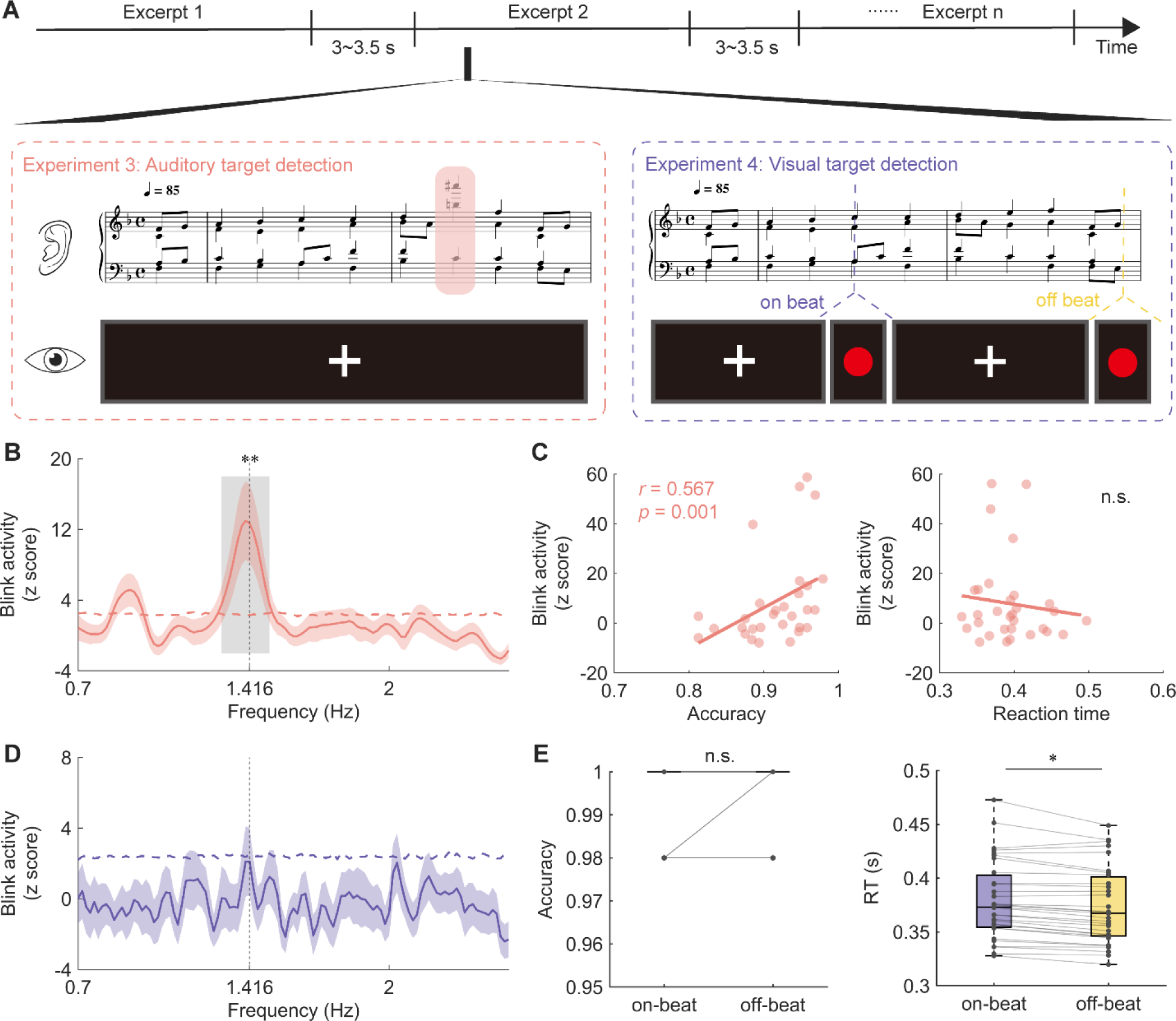
Blink tracking improves behavioral performance. (**A**) Experimental paradigm for Experiments 3-4: participants performed a pitch deviant detection task (Experiment 3) or visual target detection task (Experiment 4) while their eye movements were recorded. (**B**) Blink amplitude spectra in Experiment 3, with horizontal dashed line representing surrogate test threshold (*p* < 0.01). Shaded area represents 1 SEM across participants (n = 31). Gray box indicates frequency range above the threshold. The blink activity showed a salient peak around the beat rate. (**C**) Correlation between blink activity and behavioral indexes in Experiment 3. Colored dots indicate individuals. Blink activity was positively correlated with detection accuracy, but not with reaction time (n.s., not significant). (**D**) Blink amplitude spectra in Experiment 4 (n = 32), with no significant peak. (**E**) Group-averaged accuracy and reaction time in on-beat (purple) and off-beat (yellow) conditions. Colored connecting lines represent individual participants. Participants responded significantly faster to off-beat visual target than to on-beat target. * *p* < 0.05.

We identified the same signature of eye tracking of beats (**Fig. 6B**). We further found a positive correlation between the strength of eye tapping and the deviant detection accuracy (*r*_(30)_ = 0.567, *p* = 0.001; **Fig. 6C** left). However, no significant association was observed between eye tapping and reaction time (*r*_(31)_ = −0.072, *p* = 0.701; **Fig. 6C** right). These results suggest that proficient eye tapping may facilitate the detection of deviant pitch by aligning temporal attention with beat onset, implying a potential role of eye tapping in a dynamic attending process.

### Eye tapping indicates cross-modal interaction in music listening

Lastly, as Experiment 3 indicates that eye tapping reflects an underlying dynamic attending process, and considering one of the functions of eye blinks is to regulate the flow of visual information, we asked whether musical rhythms induce eye tapping and subsequently guide visual sensory sampling, a cross-modal process of dynamic attending.

The experimental procedure mirrored that of Experiment 3, except that 32 non-musician participants were required to detect a simply visual target—a red dot—presented on the screen while listening to music (**Fig. 6A**). The deliberate simplicity of the visual task was intended to facilitate participants’ attention towards the musical stimuli. We manipulated the temporal relationship between the visual target and musical beat onsets to explore auditory-visual interaction, resulting in two conditions: an on-beat condition and an off-beat condition. If the temporal signature of eye tapping indeed reflects that the modulation of visual sampling efficiency by musical rhythms, we would expect to observe better performance, like shorter reaction time, in the on-beat condition than the off-beat condition.

We repeated the analysis procedures on eye blinks as in Experiment 1-3, yet eye tracking of beats did not yield significant results (**Fig. 6D**). This observation is interesting, particularly considering that participants were not heavily engaged in the visual task, evidenced by a ceiling effect in the task performance with nearly 100% accuracy in both conditions (*t*_(27)_ = −0.812, *p* = 0.424, Cohen’s *d* = −0.153, **Fig. 6E** left). Our interpretation suggests that eye tapping is a spontaneous behavior induced by active engagement in music listening.

We further compared reaction time for detecting the visual target between the on-beat and off-beat conditions using a two-sided paired t-test. The result showed significantly slower responses to the on-beat visual target compared to the off-beat visual target (*t*_(30)_ = 2.417, *p* = 0.022, Cohen’s *d* = 0.434, **Fig. 6E** right). In sum, results from Experiment 4 demonstrates a cross-modal temporal modulation of visual sensory sampling during music listening.

## Discussion

We’ve shown through a sequence of studies that peripheral oculomotor activity, specifically eye blinking, consistently aligns with regular musical beats—a phenomenon we call ‘eye tapping’, as well as its neural substrates and functional roles.

Our study was initially designed to explore whether eye blinks corresponded with musical phrasal structures in our musical materials over a span of 5 to10 seconds, given the spontaneous blinking rate falls within this timeframe (12 to 15 times per minute)(*46*). Unexpectedly, as depicted in **Fig. 2**, we discovered that eye blinks tracked musical beats at 85 beats per minute and selectively adapted to different phases of musical beats. Moreover, we observed a correspondence between eye tracking and neural tracking of beats (**Fig. 3**). How is such the synchronization accomplished on the structural level? We further demonstrated that the symmetry of posterior segment of SLF had an important impact on the strength of eye tapping (**Fig. 4**). We then replicated this eye-tapping phenomenon across a broader range of tempi (66 – 120 beats per minute) and in auditory sequences devoid of melodic cues (**Fig. 5**), affirming its robustness. Importantly, our investigations into the functional relevance of eye tapping revealed intriguing results. Experiment 3 revealed a correlation between better eye tapping and improved auditory perception in a music-dynamic environment, while Experiment 4 highlighted the cross-modal impact of music on visual sampling (**Fig. 6**).

Eye activities are vital for identifying mental processes and formulating hypotheses in cognitive and neural function studies(*54*). Recent research has shed light on the intricate relationship between eye movements and music listening(*28, 29*), aiming to uncover the cognitive complexities of human music activities related to emotion, memory, anticipation, and attention. However, a direct connection between eye movements and beat structures remains elusive. While typical voluntary actions, such as finger tapping or dancing, have been employed to study musical rhythm perception, an intriguing question that remains unanswered is whether music can elicit spontaneous autonomic activities. Our discovery of eye tapping establishes a robust link between eye blinks and rhythmic musical structures in a naturalistic listening setting, demonstrating a novel spontaneous music-synchronizing behavior. This is particularly interesting because, when people tap their feet or nod their heads “involuntary”, they are aware of doing so. However, although we did not measure participants’ self-awareness of their blinking behavior in the current study, no participant reported being aware of blinking along with the beat after the experiment. The fact that autonomic oculomotor activity can track musical beat may actually reflect the evolution-rooted primitive instinct in music rhythm processing(*20*).

In other listening context, eye blinks have also been studied. For instance, in spoken language processing, findings akin to ‘eye tapping’ exist but differ mechanistically. Research indicates that eye blinks can signal artificial sentence structures, somewhat resembling high-level musical structures but not musical beats(*40*). While musical beats form the foundation, linguistic constructs reflect cognitive load dynamics, distinguishing them from ‘eye tapping’ in music(*55*). Recent studies demonstrate that eye movements align with speech dynamics and listening effort(*56*). However, due to the inherent irregularity of speech compared to musical beats, eye movements do not always predict speech dynamics, underscoring a distinct difference from ‘eye tapping’ in music.

Notably, the observed blink inhibition a few hundreds of milliseconds prior to beat onset and a surge in blinking activity after beat onset diverge from the negative mean asynchrony observed in finger tapping studies, where finger taps slightly precede beat timing(*12*), indicating an anticipation of beat timing. One plausible explanation for this disparity is that eye blinks may be strategically timed to avoid critical junctures, notably the period immediately preceding beat onset. Consistently with prior research on blink inhibition(*50, 57–59*), our findings suggest that the brain, anticipating the impending beat onset—an event of temporal significance—proactively inhibits eye blinks to prevent sensory input obstruction. Overall, an inherent anticipatory mechanism seems to underlie blink inhibition preceding beat onset, followed by a heightened blinking post-onset.

The identification of novel cognitive and neural mechanisms often starts from pinpointing unique behavioral patterns(*7*). We posit that ‘eye tapping’, a newly discovered behavior we added to the auditory-motor synchronizing repertoire, provides fresh insights on music listening and cross-modal interaction, specifically, cross-modal active sensing. Why do eye blinks ‘tap to’ beats? In the visual domain, eye blinks act as gatekeepers for visual sampling. The temporal pattern of involuntary eye blinks shows suppression before beat onset and enhancement immediately following beat onset (Fig. 2), suggesting a selective temporal visual sampling process induced by musical rhythms. This blink behavior mirrors the dynamic attending theory in auditory processing(*60, 61*), where specific acoustic patterns guide attention for efficient auditory sampling. Thus, ‘eye tapping’ may signify an involuntary cross-modal interaction between visual and auditory systems, guiding visual sampling and serving as a behavioral marker for active sensing. The observed delay in responding to the on-beat visual target, despite its ease of detection in Experiment 4 indeed provides evidence supporting this conjecture. Additionally, oscillatory enhanced attentional dominance on the auditory stream at on-beat time points may also contribute to the observed deceleration in response to on-beat visual target. One natural follow-up question is how various musical rhythms influence visual sampling and attentional allocation, as indicated by ‘eye tapping’.

Interestingly, our participants could be clustered into high synchronizers or low synchronizers based on their eye tapping ability to musical beats, which aligns with prior research showing individual differences in auditory-motor synchronization with speech or tone stimuli using vocal tract or hands effectors (*51, 52*). Moreover, we found that the microstructural integrity (indexed by FA) of the posterior segment of SLF, a tract connecting auditory and parietal regions, showed bilateral symmetry in higher synchronizers but left lateralization in low synchronizers, with stronger symmetry or right-lateralization correlating with better eye tapping synchronization. This result parallels previous finding that FA-based left lateralization of a similar tract supports higher auditory-motor synchrony with speech signals (*51*), underscoring the crucial role of microstrucutual differences in white matter pathways linking auditory and parietal motor regions in influencing individual variations in auditory-motor synchronization across auditory stimuli and motor effectors.

Musical beat tracking may also engage a subcortical-cortical interaction between the basal ganglia and premotor areas (*2, 18*). While a few studies have linked the premotor areas to periphery eye movements in non-human animals(*62, 63*), a series of studies on eyeblink conditioning have identified a close relationship between the basal ganglia and eye blinks(*64–66*). Thus, it’s plausible that the basal ganglia, stimulated by musical beats, signals frontal eye field and the oculomotor muscles, prompting eye blinks via a subcortical-to-peripheral pathway. This echoes the finding shown in **Fig. 3**, where neural tracking of beats predicted eye blinking activities. Consequently, we propose a new auditory-visual interaction pathway mediated by subcortical motor areas for future investigation.

The discovery of ‘eye tapping’ marks a significant stride in our comprehension of auditory-motor synchronization, yet the current study has certain limitations that present avenues for future exploration. Firstly, the demographic profile of participants, primarily young non-musicians with normal hearing, indicates the need to expand the sample to broader populations. Including musicians can reveal how extensive musical training influences the phenomenon, while diverse age groups could help assess developmental trajectories and the consistency of ‘eye tapping’ across the lifespan. Given that impaired rhythm processing skills may be a risk factor for developing various neurodevelopmental disorders(*6*), ‘eye tapping’ and ‘blink inhibition’(*50*) during music listening have great potential to serve as more implicit and feasible diagnostic and interventional biomarkers for children, compared to explicit paradigms like finger tapping. Secondly, the exclusive use of Bach’s chorales, noted for their high temporal regularity and representation of Western classical music, suggests a limitation in musical diversity. Investigating ‘eye tapping’ in response to music featuring irregular rhythms, syncopation, and a broad spectrum of genres and cultural backgrounds is crucial for understanding the universality of this phenomenon and its dependency on rhythmic structures. Furthermore, the coordination between blinks and pupil size dynamics in tracking musical beat structure warrants further investigation to advance our understanding of eye movements in music listening. Future research should also delve into the neural underpinnings of ‘eye tapping’, particularly focusing on the roles of the basal ganglia, frontal eye field and premotor areas. Such studies could not only enrich our knowledge of the neural mechanisms driving ‘eye tapping’ but also to explore its applications in fields like neurorehabilitation, thereby enhancing our understanding of sensory integration and motor coordination across different populations and musical contexts.

In conclusion, our study elucidates the relationship between music listening and peripheral oculomotor activity, specifically the correlation between eye blinks and musical rhythms—termed ‘eye tapping’. Our research establishes eye tapping as a distinctive spontaneous music-synchronizing behavior, and reveals its neurophysiological and structural correlates. By connecting dynamic attention and visual sampling to musical rhythm, we gain deeper insights into the coordination of auditory, motor, and visual systems, thereby enriching our knowledge of cross-modal active sensing and embodied musical perception.

## Materials and Methods

### Participants

A total of 123 young adults (70 women; mean age, 22.67 ± 2.98 years, ranging from 18 to 34 years old) with normal hearing (thresholds ≤ 20 dB SPL from 125 to 8000 Hz) took part in the study. Thirty individuals participated in Experiment 1, 30 in Experiment 2, 31 in Experiment 3 and 32 in Experiment 4. All participants were nonmusicians and scored an average of 17.45 (SD = ± 7.60, range: 7 - 39) on the musical training subscale of the Goldsmiths Musical Sophistication Index (Gold-MSI) questionnaire(*67*). All participants reported normal or corrected-to-normal vision and an absence of neurological or psychiatric diseases. They gave written informed consent prior to experiment and received a monetary compensation of ¥ 50 per hour for their participation. The study was approved by the Ethics Committee of the Institute of Psychology, Chinese Academy of Sciences.

### Stimuli

The stimuli comprised ten musical pieces selected from the 371 four-part chorales by Johann Sebastian Bach(*68*). The original musical scores were checked manually to ensure inclusion of only quarter notes, eighth notes and sixteenth notes, while removing ties across notes to facilitate repetition. Importantly, we eliminated fermatas from the original musical pieces to prompt participants to parse the musical pieces by structural cues rather than rhythmical or acoustic cues for phrasal structures (such as pauses between phrases and lengthened notes). The duration of the musical pieces ranged from 22.59 s to 59.30 s (39.82 ± 11.49 s, mean ± SD).

In Experiment 1, we created a reverse version by completely reversing the order of beats in each original piece (for an example piece, see **Fig. 1B**). Therefore, the basic musical contents were equal in both versions, but the harmonic progressions were manipulated in the reverse version which would affect listeners’ expectation or familiarity of the musical pieces.

For the rhythm condition in Experiment 2, the stimuli consisted of frequent standard tones (440 Hz) with timing structures identical to the original pieces. To ensure participants’ attention remained focused, we randomly inserted one to three chords deviating in timbre (guitar) into two additional pieces selected from the musical stimuli. These two pieces containing timbre deviants were excluded from further analyses.

In Experiment 3, we extracted twenty-four melodic sequences, each comprising two phrases, from the original pieces used in Experiment 1. Each phrase consisted of eight chords. A pitch deviant note occurred randomly within the second phrase of each melody with equal probability. To adjust difficulty levels, we created pitch deviants by shifting the melody in the upper two voice parts (soprano and alto) upwards by one to two octaves. Each sequence was manipulated into two versions with the deviant inserted in different position.

Experiment 4 utilized visual stimuli, with the target being a red circle with a diameter of 50 pixels (RGB: 255, 0, 0), displayed at the center on a black background (RGB: 0, 0, 0).

All stimuli were created and exported as wav files with a piano timbre using the MuseScore 3 software. In Experiments 1, 3 and 4, musical stimuli were played at a tempo of 85 beats per minute, while Experiment 2 employed three tempi (66, 85 and 120 beats per minute). Therefore, the note rate for each musical piece at a tempo of 85 beats per minute was around 2.833 Hz, the beat rate was around 1.416 Hz, and the phrase rate was around 0.177 Hz. Musical pieces were presented at a sampling rate of 44100 Hz, and the intensity level of each piece was normalized to 70 dB SPL.

### Experimental design and procedure

Experiment 1: The twenty musical pieces (10 pieces × 2 versions) were presented in 4 blocks, each containing 5 pieces. In each block, participants listened to each of the 5 pieces three times in a randomized order. Throughout the experiment, participants were required to fixate on a white fixation cross at the center of a black screen. After the third presentation of each musical piece, participants pressed keys to rate how much they liked it using a 5-point Likert scale, with 1 indicating low preference and 5 indicating high preference. After providing their rating, the next musical piece commenced after a delay of 3∼3.5 s (see **Fig. 1A**). Each block lasted around 10 minutes and participants allowed to rest between blocks.

Experiment 2: The melody condition and the rhythm condition were presented in separate blocks, with the conditions alternating every 5 trials. Stimuli were presented to participants according to a balanced Latin square design for every condition, and the order of the two conditions was counterbalanced across participants. Before the main experiment, participants underwent a 3-min rest period while looking at a blank screen to obtain a baseline. During each block, participants were instructed to focus on the fixation cross displayed on the screen and press the spacebar as soon as possible upon detecting a timbre deviant. Participants received training until they achieved an accuracy level of 80%.

Experiment 3: Participants were presented with two versions of the twenty-four melodic sequences, each presented twice in a pseudorandomized order, ensuring that no sequence or version was repeated consecutively. Participants were asked to carefully listen to the sequences and press the spacebar as soon as they detected a pitch deviant, while maintaining fixation on the central cross on the screen. Before the experiment, participants underwent a training session, concluding once they achieved an accuracy score above 80% on the detection task.

Experiment 4: Participants engaged in a visual target detection task while ten musical pieces, used as the original version in Experiment 1, were presented twice in a pseudorandomized order. Concurrently, the fixation cross at the center of the screen occasionally transformed into a red circle. The visual target appeared either synchronously with the beat onset (on-beat condition) or randomly at one of nine temporal positions between two beats (off-beat condition). It was ensured that 1) each musical piece contained at least one target and up to three for each condition with the number of targets per condition was the same, and 2) no piece was presented twice consecutively with visual targets occurring at different positions for the same piece. Participants were instructed to observe the fixation cross while the auditory stimuli played, and press the spacebar as soon as possible once sighting the red circle on the screen. A training session was given before the experiment until participants achieved a target detection accuracy exceeding 80%.

At the end of each experiment, participants completed the Gold-MSI questionnaire and MET to measure their musical ability. The English version of the Gold-MSI questionnaire was translated into Chinese by the experimenters. Participants in Experiment 4 did not complete the MET due to the duration of the task.

### Data acquisition

Eye-tracking data were recorded in all experiments using an Eyelink Portable Duo system (SR Research, Ontario, Canada), with a sampling rate of 500 Hz. Participants maintained a consistent eye-to-monitor distance of 95 cm, with the eye tracker positioned approximately 60 cm from their eyes. Before each experiment, a nine-point standard calibration and validation test was performed. Then a fixation cross appeared on the monitor with a resolution of 1,920 × 1,080 pixels. A re-calibration procedure was applied after each break to ensure accuracy and consistency.

In Experiment 1, EEG data were recorded using a 64-channel Quik-Cap based on the international 10–20 system (Neuroscan, VA, USA), with a sampling rate of 1000 Hz. The EEG was referenced to the average of all electrodes, with skin/electrode impedance maintained below 5 kΩ. Additionally, four electrodes were used to record electrooculography (EOG): two electrodes were placed at the outer canthus of each eye, while another two electrodes were placed above and below the left eye to record the horizontal and vertical EOG signals, respectively. Additionally, DWI data were collected from the same cohort using a 3.0 T MRI system (Simens Magnetom Prisma) with a 20-channel head coil with following parameters: repetition time (TR) = 4000 ms, echo time (TE) = 79 ms, field of view (FOV) = 192 × 192 mm^2^, voxel size = 1.5 × 1.5 × 1.5 mm^3^, diffusion-weighted gradient directions = 64, b-values = 1000 s/mm^2^ and 2000 s/mm^2^, b0 non-weighted images = 5.

### Data preprocessing

Preprocessing of eye-tracking and EEG data were performed in MATLAB (The MathWorks, Inc., Natick, MA) using the Fieldtrip toolbox(*69*) and custom scripts.

For eye-tracking data, blink detection was the primary event of interest. Blinks were identified by the Eyelink tracker’s on-line parser, which flagged instances where the pupil in the camera image was absent or severely distorted by eyelid occlusion. The time series were then divided into epochs corresponding to the duration of each musical or pure-tone sequence, aligning with the experimental trials. In Experiment 2, baseline data was derived from three continuous trials, each lasting 60 s. Within each trial, data were represented as binary strings, with values set to 1 during blink occurrences and 0 otherwise. Blinks shorter than 50 ms or longer than 500 ms were excluded from the analysis. Subsequently, mean blink duration and blink rate were calculated for each participant. Blinks with mean durations exceeding three standard deviations from the data’s mean, and trials with blink rates exceeding three times the standard deviation, were removed. Following preprocessing, eye-tracking data were down-sampled to 100 Hz for further analysis.

For continuous EEG data, single trials were extracted spanning 3 s before stimulus onset to 3 s after stimulus offset. All data were band-pass filtered between 0.7 and 35 Hz using a phase-preserving two-pass fourth-order Butterworth filter, supplemented with a 48- to 52-Hz band-stop filter to eliminate line noise. Then the data were down-sampled to 100 Hz to match the sampling rate of the eye-tracking data. An independent component analysis(*70*) with 30 principal components, implemented in Fieldtrip, was performed to remove EOG and electrocardiography (ECG) artifacts. Finally, we concatenated all trials into one matrix to derive the most common component of the EEG signals reflecting neural responses to musical pieces, using MCCA(*71*). The first MCCA component was projected back to each participant and each trial, and used for subsequent analysis.

For DWI data, MRtrix3 and FSL software(*72, 73*) were employed to preprocess the raw data. Preprocessing steps included denoising, reducing the ringing artifacts, eddy current and motion correction, and bias field correction using the N4 algorithm provided in Advanced Normalization Tools. Gradient directions were also corrected after eddy current and motion correction.

### Music amplitude modulation spectrum

To derive the amplitude envelope of the musical stimuli in Experiment 1, we constructed 64 logarithmically spaced cochlear bands between 50 to 4000 Hz using a Gammatone filterbank. The amplitude envelope of each band was then extracted from the acoustic waveform using the Hilbert transform and down-sampled to 100 Hz. For each musical piece, the amplitude envelopes of the 64 bands were averaged, and the modulation spectrum of each piece was calculated using the FFT with zero-padding of 8000 points, followed by taking the absolute value of each frequency point. Next, we normalized the modulation spectrum of each piece by dividing it by the norm of its modulation spectrum. Lastly, we estimated the mean amplitude modulation spectrum across the 10 pieces for each condition and showed them in **Fig. 1C**.

### Beat/Note tracking

To access whether blink and neural signals could track hierarchical structures in auditory sequences, we used a frequency domain analysis. The analysis window for blink data varied depending on the task. In summary, we analyzed the time series from 1 s after stimulus onset to 1 s before stimulus offset in Experiments 1 and 2, and we selected the time series from 1 s after stimulus onset to the onset of the first target in Experiments 3 and 4. The blink data for each trial and participant were then transformed into the frequency domain using a Fast Fourier Transform with zero-padding of 6000 points. Given the variable trial lengths, we averaged the amplitude spectra across the 10 pieces for each version and repetition in the frequency domain. Next, we identified the lower and upper bounds on the significant frequency ranges among all independent variables showing robust blink and neural tracking of musical structures. We then averaged the amplitude spectra within the frequency range for each variable for further investigation. For the baseline in Experiment 2, we first averaged the data across three trials in the temporal domain and then applied the same procedure for conversion to the frequency domain.

### Blink rate

In this analysis, we derived blink time series within a window from −353 to 353 ms relative to the beat onset for each beat, determined by the tempo. The blink rate at each time point was defined as the average value across trials for each participant and version. Then we identified the time points corresponding to the highest and lowest values (using the ‘findpeaks’ function from MATLAB), divided them by the beat length, and then multiplied them by 2π to obtain blink phase angles in radians for each participant for further analysis.

### Mutual information

In the above frequency-domain analysis, we observed robust blink tracking of musical sequences primarily centered around the beat rate. To estimate the statistical dependency between the neural response and the eye-tracking data, we focused on the EEG data at the beat rate and computed mutual information (MI) between it and the blink signal. First, we extracted the time-frequency distribution of EEG data at the beat rate using a Morlet wavelet (with a sliding window length of 3) implemented in Fieldtrip, with steps of 10 ms. Next, we converted the output, corresponding to the EEG power, to a decibel scale using a pre-stimulus baseline ranging from −1500 to −500 ms. Then, MI between the blink signal and EEG power was calculated using Gaussian Copula Mutual Information(*74*) at different lags: the blink signal was shifted against the EEG power from −200 to 200ms in 20-ms steps. Finally, we summed the MI values across lags and averaged them across 10 pieces for each version and repetition for further analysis.

### Temporal response function

What’s the relationship between ocular activity and neural response in the time domain? To reveal the temporal dynamics of the neural response to blink activity, we employed the mTRF toolbox(*75*) to estimate the temporal response function (TRF) of EEG power at the beat rate, utilizing the blink onset vector as the input stimulus feature. The TRF, denoted as w, was estimated using the following equation (Eq. 1):

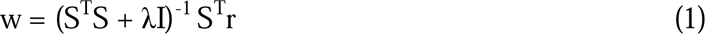

Here, S is the lagged time series of the input stimulus feature (the blink onset vector obtained from the eye-tracking data), r is the continuous neural response at the beat rate, I is the identity matrix, and λ is a constant ridge parameter.

The TRF for each musical piece and participant was calculated using the blink onset vector and EEG power from 1 s after stimulus onset to 1 s before stimulus offset. The time lags ranged from 2500 ms before blink onset to 2500 ms after, with λ set to 0. Then we averaged the TRF weights across 10 pieces for each version and repetition for further permutation test (described in the following section), and extracted the TRF weights at the peak time points for all independent variables for further comparison.

### Fiber tractography

Fiber tractography based on high angular resolution diffusion imaging (HARDI) was performed using MRtrix3. Firstly, 3-tissue (white matter, gray matter, and cerebrospinal fluid) response functions were obtained by the command ‘dwi2response dhollander’(*76*). Secondly, fiber orientation distributions (FOD) of each voxel were estimated using multi-shell multi-tissue constrained spherical deconvolution algorithm(*77*). Thirdly, the whole-brain probabilistic tractography was performed to generate ten million streamlines for each participant based on iFOD2 tracking algorithm(*78*). Lastly, track filtering was performed to attain 1 million streamlines(*79*).

The target two segments of the SLF were defined according to a simple classification system(*80*): the dSLF, namely the classical SLF II, connects the Geschwind’s area and the dorsolateral frontal area; the pSLF corresponding to the SLF-tp or posterior AF connects the Geschwind’s area and the Wernicke’s area. Three ROIs (Geschwind’s area: super marginal gyrus (SMG) and angular gyrus (AG); dorsolateral frontal area: posterior superior frontal gyrus (SFG) and middle frontal gyrus (MFG); Wernicke’s area: posterior superior temporal gyrus (STG) and middle temporal gyrus (MTG)) were extracted from individual anatomical image parceled by Freesurfer for each hemisphere to dissect bilateral segments of SLF. **Supplementary Fig. 3** shows fiber tractography of the dorsal and posterior SLFs in bilateral hemispheres from one participant.

We extracted three indexes to measure the microstructural properties of fibers: FA of diffusion tensor imaging (DTI) as well as NDI and ODI derived from neurite orientation segment were extracted for each participant. LI for each parameter was subsequently calculated using the following equation (Eq. 2):

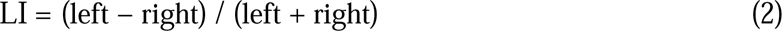

### Behavioral measure

Accuracy and reaction time were recorded in Experiments 3 and 4. Detection accuracy was defined as the ratio of correctly detected targets within one beat duration to the total number of targets presented. Only correct trials were included in the reaction time analysis.

### Statistical analysis

All statistical analyses were calculated using MATLAB and SPSS 20.0 (IBM Corp., Armonk, N.Y., USA). Group statistical analyses were performed on the data of all participants. Prior to analysis, outliers were identified and removed using the mean detection method.

For behavioral data, statistical differences between conditions (**Fig. 1D and 6E**) were evaluated using a paired t-test with a threshold of *p* = 0.05. For the ocular and neural activities, all statistical analyses (**Fig. 2B, 3C, 3E, 5D** left and **Supplementary Fig. 2B**) were performed with repeated-measures ANOVA to examine the effects of reversal manipulation, stimulus repetition, stimulus type and tempo. To address the issue of multiple comparisons, the statistical significance level was set at a corrected *p* < 0.05 using the Bonferroni correction method. The Greenhouse-Geisser correction was used when the assumption of sphericity was violated. The individual differences in blink response amplitude (**Fig. 4A**) were evaluated by a k-means clustering method(*82*) with two clusters and five replicates using squared Euclidean distance. Statistical differences between groups (**Fig. 4B and 4E**) were evaluated using an independent t-test with a threshold of *p* = 0.05. For neural activities, group comparisons (**Fig. 4C**) were evaluated using a Mann-Whitney U test. All statistical tests performed were two-tailed. Correlations between blink/neural response and behavioral indexes (**Fig. 2C, 5D** right, **6C** and **Supplementary Fig. 2C**) were estimated using Spearman’s rank correlation coefficients and correlations between blink response and microstructural literality indexes (**Fig. 4F**) were estimated using Pearson’s correlation coefficients according to the data distribution.

To identify the significant frequency ranges showing blink/neural tracking of auditory sequences (**Fig. 2A, 3A, 5B, 5C, 6B and 6D**), we completed a surrogate test. Blink/neural data underwent the same Fourier transform processing steps described above, yielding a complex number at each frequency point for each trial and participant. Then the data were phase shuffled in the frequency domain by multiplying the data with a complex number which has a magnitude of 1 and a randomly generated phase between 0 and 2π. As for the frequency domain analysis, we averaged the amplitude spectra of surrogate data across trials for each condition. This procedure was repeated 1000 times, creating a null distribution of 1000 surrogate amplitude modulation spectra for each condition, from which the 99^th^ percentile was chosen as the significant threshold. Finally, we z-scored the raw modulation spectra with respect to the mean of the surrogate distribution for each condition. For baseline data (**Fig. 5B**), the surrogate test procedure was identical, except for data surrogation in the temporal domain.

To reveal the change trend of blink rate throughout the experimental period (**Fig. 2D**), we used a permutation test: the blink raw data for each participant and each trial was circularly shifted by a random shift amount within the time series, and blink rate was recomputed from the new data. We repeated this process 1000 times and derived the first and 99th percentiles of permuted blink data to identify significant increase and decrease in blink rate. The distribution of blink phase angles was assessed by the V test with a threshold of *p* = 0.05.

To assess the statistical significance of the coupling between the blink signal and the beat-rate EEG response (**Fig. 3B**), we conducted a permutation test. Specifically, the beat-rate EEG power for each participant and trial was circularly shifted by a random shift amount within the time series, and MI between the time-shifted data and blink signal was recalculated. This procedure was repeated 1000 times to create a null distribution of surrogate MI values for each version and repetition, from which a p-value was obtained by counting how many surrogate MI values exceeding the empirical MI value.

To confirm that the rise in TRF weights before blink onset was attributable to pre-onset prediction for the beat-rate neural response to blink (**Fig. 3D**), a permutation test was completed following the same procedure used for **Fig. 2D**. The TRF was estimated between the new blink signal and EEG power 1000 times, and a threshold of a one-sided alpha level of 0.01 was derived for statistical significance.

## Acknowledgments

We thank Lidongsheng Xing and Jingwen Wang for their assistance with data collection; Pauline Larrouy-Maestri and Hou Chen for their assistance in preparing music materials; Baishen Liang, Xiang Li, Lei Zhang, Lyu Baihan and Peiqing Jin for their help with data processing. We thank Robert Zatorre for his comments on a previous version of the manuscript.

## Funding

Strategic Priority Research Program of Chinese Academy of Sciences XDB32010300 (Y.D.) STI 2030—Major Projects 2021ZD0201500 (Y.D.)

Improvement on Competitiveness in Hiring New Faculties Funding Scheme, the Chinese University of Hong Kong (4937113, X.T.).

## Author contributions

Conceptualization: X.T., Y.D.

Methodology: Y.W., X.T., Y.D.

Investigation: Y.W.

Visualization: Y.W.

Funding acquisition: X.T., Y.D.

Writing – original draft: Y.W., X.T.

Writing – review & editing: X.T., Y.D.

## Competing interests

Authors declare that they have no competing interests.

## Data and materials availability

All data are available in the main text or the supplementary materials. Preprocessed data that support the findings of this work are available in the Open Science Framework repository (https://osf.io/xhfn4/). Raw data will be made available upon request.

## Notes

https://osf.io/xhfn4/

### Competing Interest Statement

The authors have declared no competing interest.

### Summary of Updates

New results from DTI were added.

