## Supplementary for "Eyes robustly blink to musical beats like tapping"

Yiyang Wu *et al.*

**This PDF file includes:**

Supplementary Text

Figs. S1 to S3

Table S1

Supplementary Text

To confirm that the reversal manipulation of harmonic progressions in Experiment 1 didn’t affect liking ratings of musical pieces, we analyzed the ratings of liking for two reversal versions. As expected, there was no significant differences in liking ratings between original and reverse versions (*t*_(29)_ = 0.715, *p* = 0.480, Cohen’s *d* = 0.131; **Fig. 1D**). This finding suggests that liking would not cause the differences in blink/neural signals between the reversal versions found later.

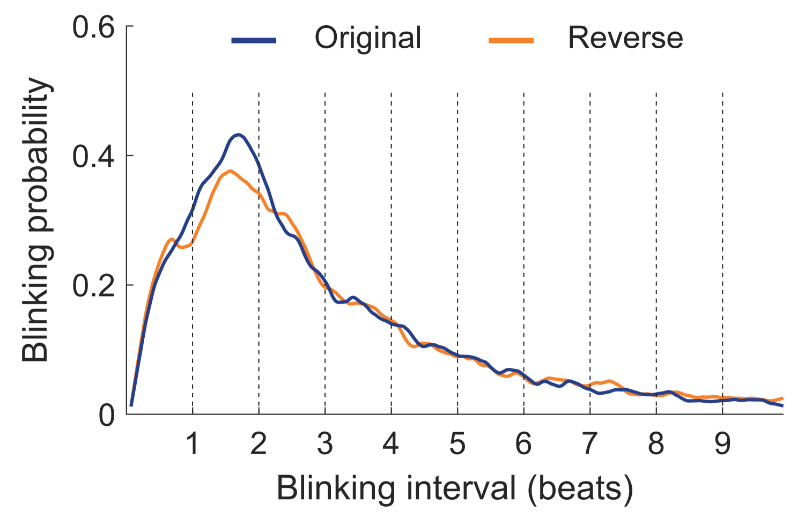

Fig. S1.

**Blink probability at different time interval for two versions.** The length of each musical beat is 706 ms. The figure displays that the inter-blink interval is most often one to two beats, consistent with what we observe in Fig. 2.

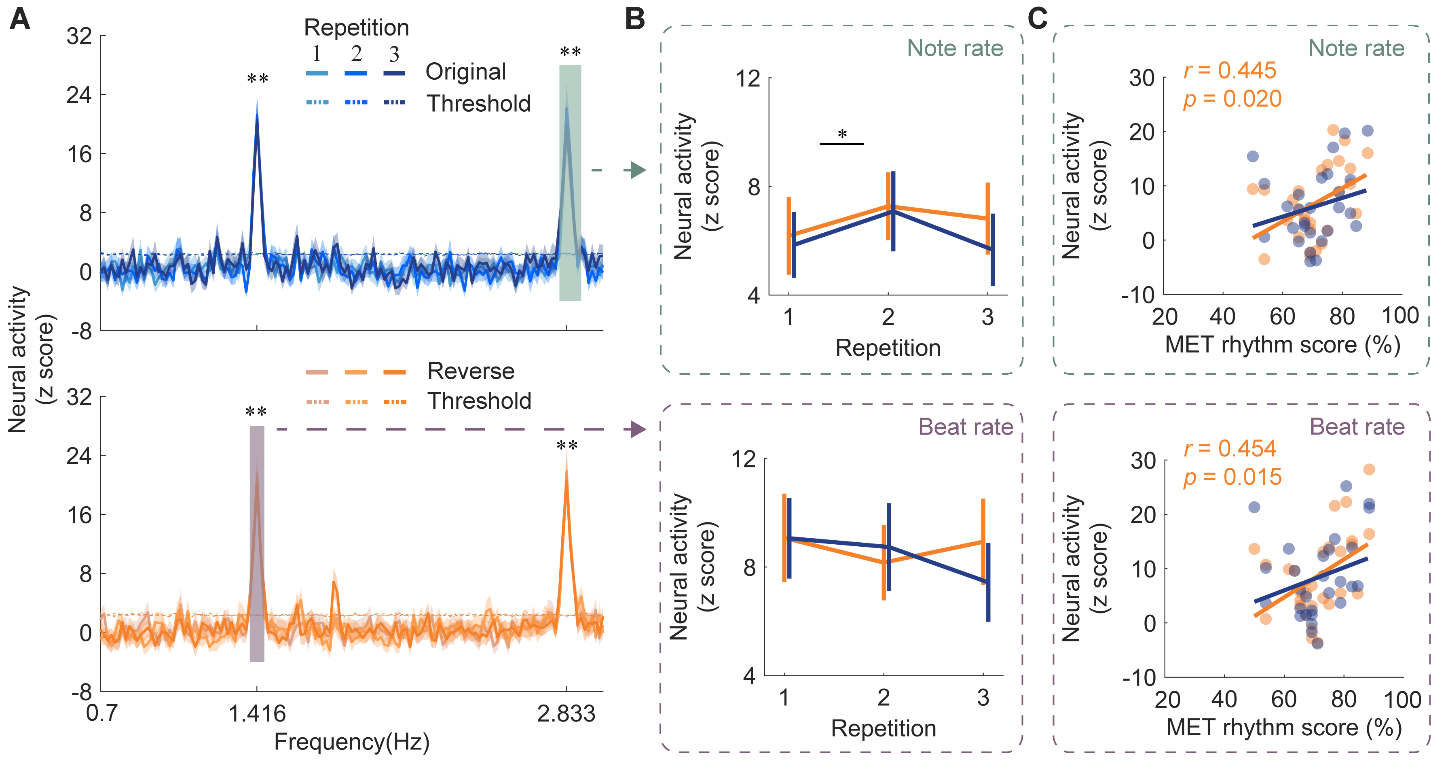

Fig. S2.

**Neural tracking of musical beats and notes.** (**A**) Neural response spectra for two versions consistent with Fig. 3A. The colored boxes indicate the frequency ranges where the amplitude was above the threshold. (**B**) EEG response amplitude at note and beat rates. Error bars denote 1 SEM across participants. Note-rate EEG response increased with repeated exposure of music. (**C**) Correlation between the MET rhythm score and EEG response amplitude at note and beat rates. The two EEG responses were both correlated with the MET rhythm score for the reverse version. Colored dots indicate individuals. * *p* < 0.05, ** *p* < 0.01.

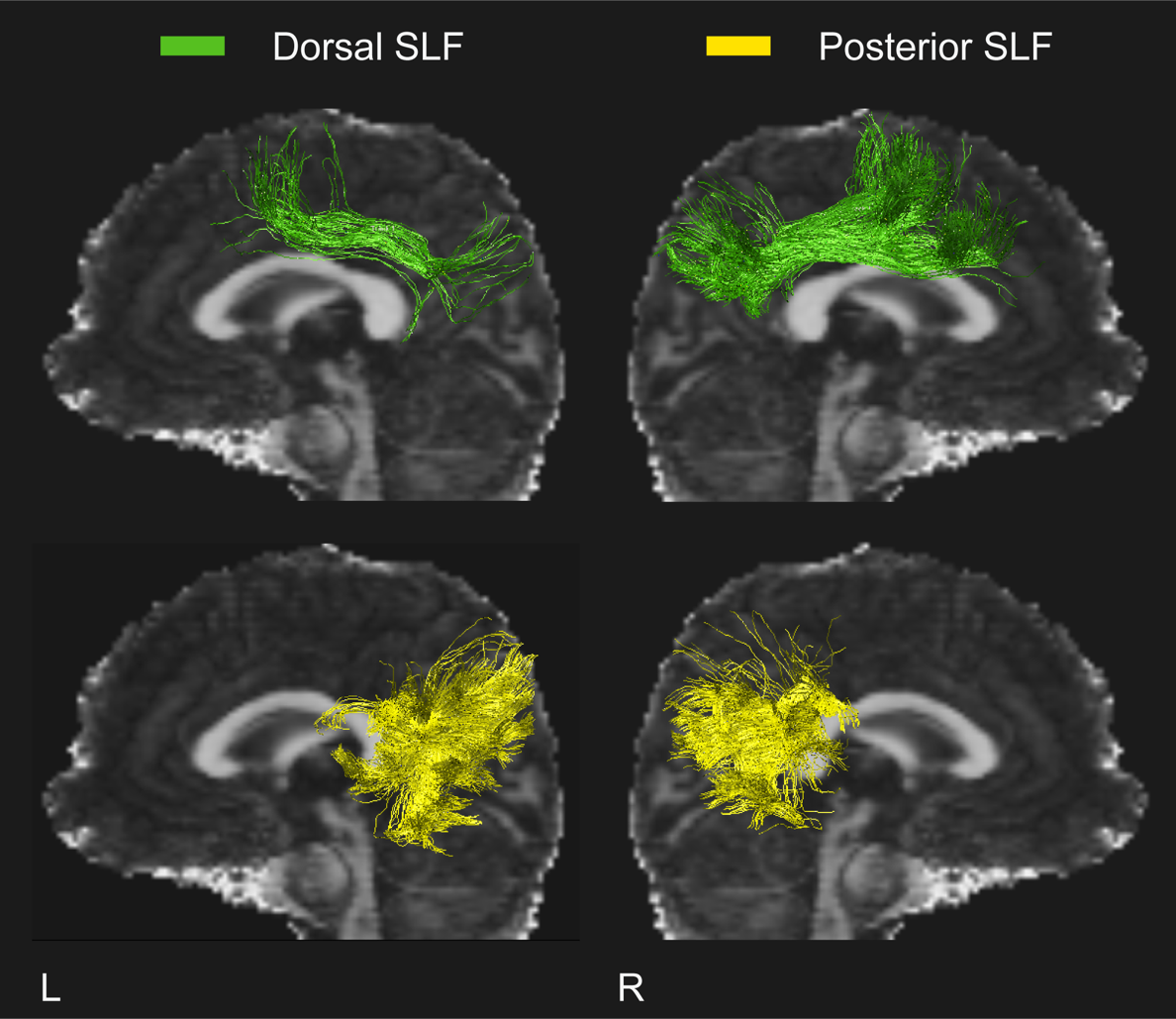

Fig. S3.

**The tractography of two SLF segments in bilateral hemispheres from a participant.** Dorsal and posterior segments of SLF are depicted in green and yellow.

Table S1.

Group comparisons of microstructural and laterality index between low synchronizers and high synchronizers.

|  | **Microstructural index** | | | **Laterality index** | | |
| --- | --- | --- | --- | --- | --- | --- |
|  | **FA** | **NDI** | **ODI** | **FA** | **NDI** | **ODI** |
| **L dorsal SLF** | -1.282  (0.212) | 0.423  (0.676) | 1.765 (0.090) | -0.829  (0.417) | 0.152 (0.881) | 0.747  (0.462) |
| **R dorsal SLF** | -0.370 (0.715) | 0.287  (0.777) | 1.112  (0.277) |  |  |  |
| **L posterior SLF** | 0.882  (0.387) | 1.500  (0.147) | -0.235 (0.816) | **2.156 (0.041) *** | 1.094  (0.285) | -1.711 (0.100) |
| **R posterior SLF** | -0.684  (0.500) | 0.490 (0.628) | 1.332  (0.195) |  |  |  |

SLF, superior longitudinal fasciculus; FA, fractional anisotropy; NDI, neurite density index; ODI, orientation dispersion index. Values are *t (p)*. *P* was estimated by two-sided independent t-test with a threshold of *p* = 0.05. * *p* < 0.05.
